# *streammd*: fast low-memory duplicate marking using a Bloom filter

**DOI:** 10.1101/2022.10.12.511997

**Authors:** Conrad Leonard

## Abstract

**Summary:** The identification of duplicate reads is an essential pre-processing step in short-read sequencing analysis. For large sequencing libraries this step is typically time-consuming and resource-intensive. Here we present `streammd`: a fast, memory-efficient, single-pass duplicate marking tool operating on the principle of a Bloom filter. We show that `streammd` closely reproduces the outputs of Picard `MarkDuplicates`, a widely-used duplicate marking program, while being substantially faster and suitable for pipelined applications, and that it requires much less memory than `SAMBLASTER`, another single-pass duplicate marking tool.

**Availability and Implementation:** `streammd` is a C++ program available from GitHub (https://github.com/delocalizer/streammd) under the MIT license. Install instructions are in the `README.md` file. Unit tests are runnable with `make check`. Open issues are listed at https://github.com/delocalizer/streammd/issues.

**Supplementary information:** Supplemenatary_figures.zip

Supplementary_tables.zip

## Introduction

Duplicate templates commonly occur in short-read sequencing (SRS) data via a number of mechanisms — PCR duplicates from library preparation, optical duplicates (sequencing artefacts) or ‘natural’ duplicates in which fragments from the same genomic coordinates are generated from different molecules by chance. It is often desirable to identify duplicate templates in SRS data, for example for library complexity estimation, or for removal from consideration for analysis when the true number of unique molecules is important, e.g. allele frequency calculation or low-frequency somatic variant calling.

From modern patterned flowcell technologies the number of duplicates due to sequencing artefacts is small. Of the remaining duplicates it is not normally possible to distinguish between PCR duplicates and natural duplicates — although, see Salzberg et al (2018) for one approach — but it is generally assumed for most library protocols that the number of natural duplicates is also small, and therefore that most duplicate templates are PCR duplicates.

With paired-end sequencing technologies duplicate identification is readily performed after alignment by recognizing as duplicates all templates whose ends are located at the same genomic coordinates. A widely-used tool for this purpose is `MarkDuplicates` from the Picard Toolkit (2019). It is also possible to make this a streaming algorithm taking as inputs the output directly from the aligner: storing the ends of each template as it is received in a hash table, and identifying subsequent matches as duplicates, e.g. `SAMBLASTER` (Faust & Hall, 2014). With a conventional hash structure the memory requirements of this approach may be considerable for large libraries — a 60x coverage human whole-genome BAM file is around 1B templates and the resulting hash structure tens of GB. In this work we instead use a Bloom filter (Bloom, 1970) to achieve fast streaming duplicate marking in a small memory footprint, even at low false-positive rates.

## Implementation

`streammd` is implemented as a C++ program that runs in a single process. A Bloom filter is initialized with k=10 hash functions and a bit array sized to meet the user-specified false-positive rate and memory allowance. Input is QNAME-grouped SAM records; paired-end reads are accepted by default but single-end reads may also be specified. Template ends are calculated from the POS and CIGAR fields of the primary alignments in an input QNAME group and the signature hashed into the Bloom filter. If the signature is detected as already present the SAM record is marked as a duplicate in the output — the 0×400 bit is set on the FLAG and the `PG:Z:streammd` tag is added. At the end of processing metrics are written to file in JSON format. By default an error is generated if the capacity of the Bloom filter is exceeded, i.e. the number of stored items exceeds that at which the false-positive rate is predicted to be reached. This behavior can be toggled with `--allow-overcapacity`.

In addition to unit tests included with the tool, to demonstrate correctness we take Picard Toolkit to be the reference implementation and compare in Figure 1a) the counts of duplicate FLAG values in the outputs from `MarkDuplicates` and `streammd`. The input consists of 4 × 10^8^ SAM records comprising 2 ×10^8^ templates from a single read group of 2×101bp Illumina paired-end sequencing. `streammd` was run with default false-positive rate of 10^-6^.

**Fig. 1.**
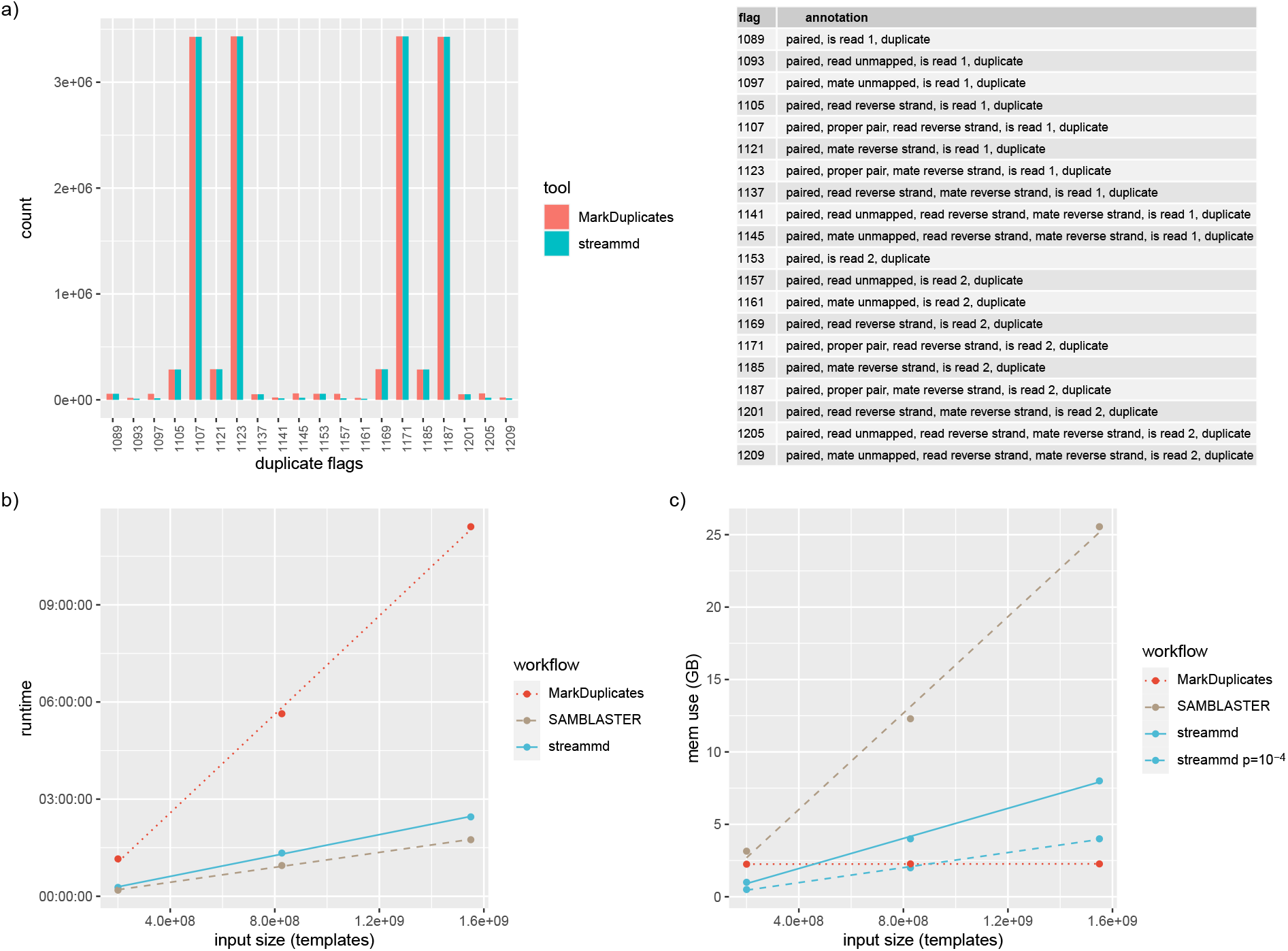
**a)** Duplicate flag counts after duplicate marking with `streammd` and Picard `MarkDuplicates`. Input size = 3.97×10^8^ reads = 1.99×10^8^ templates; `streammd` run with *p*=10^-6^. **b)** Runtime for 3 input sizes. **c)** Memory use for 3 input sizes. The small, medium and large inputs correspond to single library paired-end human WGS bams of 30G, 117G and 227G: approximately 8x, 30x and 60x read depth.

To demonstrate performance and resource use, we show in Figure 1b) and 1c) runtime and memory use respectively for `streammd`, `MarkDuplicates` and `SAMBLASTER` run on three different sized inputs. The small and medium inputs correspond to 2 × 10^8^ templates and 8 × 10^8^ templates respectively of 2×101bp Illumina paired-end sequencing of a single library prepared from human lymphocyte sample MELA_0102 (Hayward, Wilmott, Waddell, et al 2017). The large input corresponds to 1.6 × 10^9^ templates of 2×100bp BGI paired-end sequencing of a single library prepared from human melanoma cell-line sample COLO-829. These correspond to raw read depths across GRCh38 of approximately 8x, 30x and 60x respectively. Inputs were supplied to all tools in QNAME-grouped SAM format, and the output of `MarkDuplicates` specified as SAM format to avoid compression overhead. The latest versions available of all tools were used — `streammd` 4.0.2, Picard Toolkit 2.27.4, and `SAMBLASTER` 0.1.26. All tools were run alone on a 12th generation Dell c6320 with 256G RAM and dual Intel^®^ Xeon^®^ CPU E5-2690 v4 @ 2.60GHz processors. `streammd` and `SAMBLASTER` were compiled on the same platform using `g++` with `-03. MarkDuplicates` was run using Java^®^ JRE 1.8.152, 2G heap space, and serial GC. `streammd` was run with both the default false-positive rate of 10^-6^ and a more lenient 10^-4^ rate, and memory allowance of 1GiB, 4GiB and 8GiB for the small, medium, and large inputs respectively.

## Discussion

In Figure 1a) it can be seen that the results from `streammd` and `MarkDuplicates` are highly concordant. The small substantive differences are in flags involving single unmapped ends (1093, 1097, etc) due to differing logic between the implementations for identifying duplicates involving these templates. Plot and counts of all FLAG values can be found in Supplementary Figure 1 and Supplementary Table 1 respectively.

In Figure 1b) it is observed that `streammd` averages ~ 4.5 × as fast as `MarkDuplicates` at all input sizes, as we might expect for an efficient single-pass operation. From the same figure we observe that `SAMBLASTER` averages ~ 1.4× as fast as `streammd`, which is to be expected since `SAMBLASTER` performs only one hash and address operation per template, where `streammd` performs 10. Profiling with gprof reveals that the majority of the execution time in `streammd` is in fact bit array access, something of a testament to the speed of the xxhash algorithm used here (Collett, 2012).

In Figure 1c) it is observed that `streammd` uses ~ 1/3 the memory of `SAMBLASTER` at all input sizes when using the default false-positive rate of 10^-6^, and ~ 1/6 the memory of `SAMBLASTER` using a false-positive rate of 10^-4^. `MarkDuplicates` was run at all input sizes with the recommended 2G of heap (Picard Toolkit, 2019).

To complete the evaluation of resource use, in Supplementary Figure 2 we plot maximum and average CPU usage for the three tools. `SAMBLASTER` and `streammd` use a single CPU core, while `MarkDuplicates` averages 1.25 and peaks at 4 due to JVM overhead (garbage collection).

Depending on workflow requirements, potential limitations of `streammd` include its inability to pick the “best” among duplicates — a limitation of the streaming duplicate marking approach generally — and loss of some small fraction of unique sequence coverage due to Bloom filter false positives. We believe that the default setting of 10^-6^, and often higher, is tolerable in most applications. We note that a Bloom filter cannot yield false negatives; all true duplicates will be marked as such.

## Conclusion

`streammd` achieves fast, accurate duplicate marking of large sequencing libraries in a small memory footprint using the principle of a Bloom filter. Its single-pass operation and low memory use per core make it attractive for pipelined workflows, increasing the benefit of fast processing speed and allowing efficient packing of alignment and post-alignment tasks in HPC and container orchestration environments.

## Acknowledgments

Thanks to John Pearson and the Genome Informatics team for useful discussions.

## References

1. Anna C. Salzberg, Jiafen Hu, Elizabeth J. Conroy, Nancy M. Cladel, Robert M. Brucklacher, Georgina V. Bixler, Yuka Imamura Kawasawa Ph.D (2018) Effects of duplicated mapped read PCR artifacts on RNA-seq differential expression analysis based on qRNA-seq bioRxiv 301259; doi: https://doi.org/10.1101/301259

2. Picard Toolkit. 2019. Broad Institute, GitHub Repository. https://broadinstitute.github.io/picard/;BroadInstitute

3. Faust, G. G., & Hall, I. M. (2014). SAMBLASTER: fast duplicate marking and structural variant read extraction. In Bioinformatics (Vol. 30, Issue 17, pp. 2503–2505). Oxford University Press (OUP); https://doi.org/10.1093/bioinformatics/btu314

4. Bloom, B. H. (1970). Space/time trade-offs in hash coding with allowable errors. In Communications of the ACM (Vol. 13, Issue 7, pp. 422–426). Association for Computing Machinery (ACM); https://doi.org/10.1145/362686.362692

5. Hayward, N., Wilmott, J., Waddell, N. et al. Whole-genome landscapes of major melanoma subtypes. Nature 545, 175–180 (2017). https://doi.org/10.1038/nature22071

6. Yann Collett. xxHash-Extremely fast hash algorithm. GitHub https://github.com/Cyan4973/xxHash (2012-2022)

7. Broad Institute. Picard documentation https://broadinstitute.github.io/picard/command-line-overview.html#Overview (2022)

