## Supplementary figures and images for "*streammd*: fast low-memory duplicate marking using a Bloom filter"

### Supplementary_figure_1.png

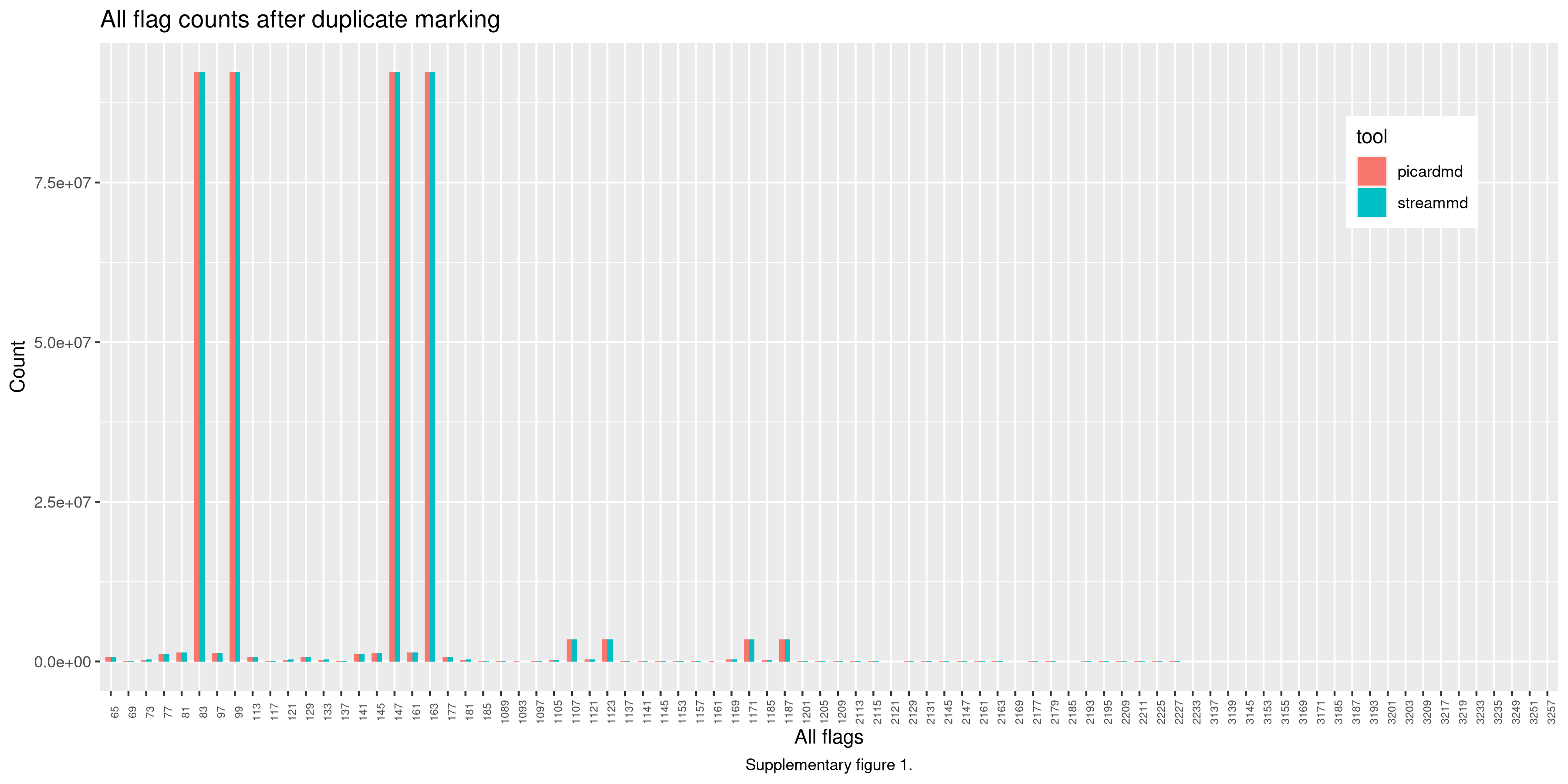

### Supplementary_figure_2.png

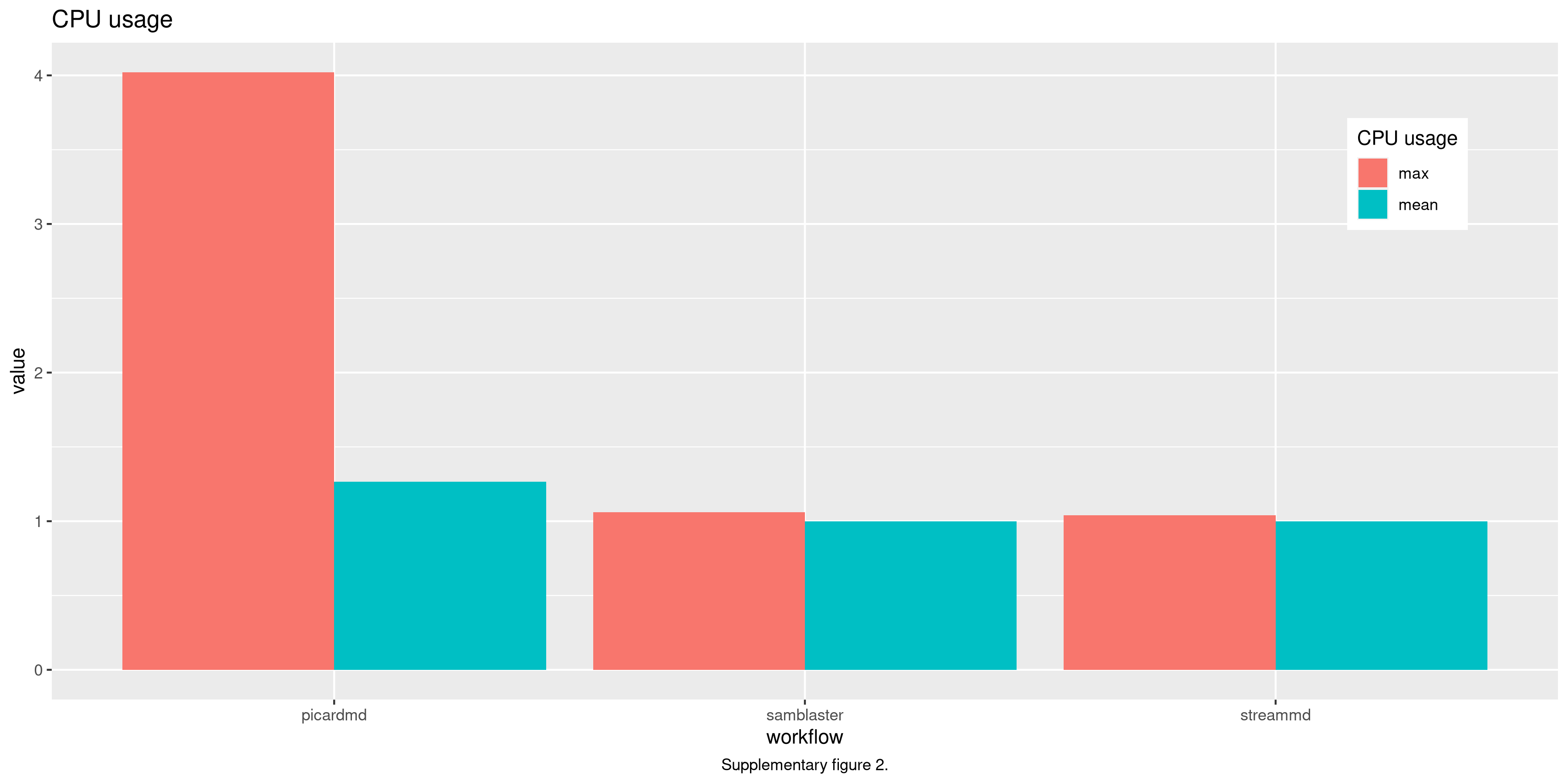
